# First detection of *Cytauxzoon* spp. DNA in questing *Ixodes ricinus* nymphs

**DOI:** 10.1101/2025.08.04.667914

**Authors:** Marina L. Meli, Theres Meili, Benita Pineroli, Eva Bönzli, Ramon M. Eichenberger, Barbara Willi, Regina Hofmann-Lehmann

## Abstract

Feline cytauxzoonosis is an emerging tick-borne disease in Europe. While infections have been reported in different European countries, the tick vector remains unknown. This study investigated 665 ticks collected in 2019 (n= 160), 2022 (n= 7) and 2024 (n= 658) in a *Cytauxzoon* spp. hotspot region in central Switzerland (62 ticks from cats; 603 ticks from vegetation). Ticks were morphologically characterized, pooled by origin and life-stage, screened for *Cytauxzoon* spp. 18S rRNA by qPCR and conventional PCR and positive samples confirmed by sequencing. All ticks belonged to *Ixodes ricinus* (50 males, 83 females, 532 nymphs). Four tick pools from 2019 tested *Cytauxzoon* spp. positive: one pool of 4 non-engorged male ticks from cats and three pools of 5-6 nymphs each from vegetation. All ticks collected in 2022 and 2024 tested negative. Amplification of the almost full-length (1535 bp, 1 pool) or partial (219-140 bp, 3 pools) 18S rRNA gene revealed a sequence identify of 98.6–100% with *Cytauxzoon* spp. previously detected in cats from this area. The detection of *Cytauxzoon* spp. in questing *Ixodes ricinus* nymphs suggests a potential role of this tick species in the parasites’ transmission cycle in Central Europe and raises the possibility of transovarial transmission. Mitochondrial gene sequencing was unsuccessful, but the detected *Cytauxzoon* spp. likely represent *Cytauxzoon europaeus*. Discrepancies between qPCR and conventional PCR results point to possible amplification of tick endosymbionts, highlighting the importance of confirmatory sequencing particularly when testing tick-derived DNA. In conclusion, this is the first report of *Cytauxzoon* spp. in questing *I. ricinus* ticks in Europe. Our findings underscore the need for further research to confirm vector competence and clarify transmission dynamics.

## Introduction

*Cytauxzoon* species are vector-borne apicomplexan hemoparasites that infect both wild and domestic felids globally. Multiple species within the genus *Cytauxzoon* have been identified. Among them, *Cytauxzoon felis* is known to cause severe and often fatal disease in domestic cats and has been well characterized in domestic and wild felids in the United States (Wang et al., 2017). The bobcat (*Lynx rufus*)—which typically experiences asymptomatic infections—is considered the natural reservoir of *C. felis* in the United States, but other wild felid species (mountain lions, cougars, ocelots, margays, and jaguars) as well as subclinically infected domestic cats, may also serve as reservoirs (Birkenheuer et al., 2008; Brown et al., 2008; Brown et al., 2010; Garner et al., 1996; Glenn et al., 1983; Meinkoth et al., 2000; Rotstein et al., 1999; Yabsley et al., 2006). In recent years, organisms closely related to *C. felis* have been reported in domestic cats in China, India and Iran, and in domestic and wild felids in Brazil (Calchi et al., 2025; Malangmei et al., 2020; Rahmati Moghaddam et al., 2020; Zou et al., 2019). Together, these “new world isolates” of *Cytauxzoon* spp. are phylogenetically different from “old world isolates” of *Cytauxzoon* spp. The latter comprise *Cytauxzoon manul* isolated from a Pallas’s cat (*Otocolobus manul*) from Mongolia (Ketz-Riley et al., 2003), and *Cytauxzoon* spp. isolated from domestic and wild cats in different regions in Europe (Alho et al., 2016; Antognoni et al., 2022; Carli et al., 2014; Diaz-Reganon et al., 2017; Grillini et al., 2021; Legroux et al., 2017; Naidenko et al., 2022; Nentwig et al., 2018; Panait et al., 2020; Spada et al., 2014; Veronesi et al., 2016; Willi et al., 2022). Most recently, new species of *Cytauxzoon* spp., *Cytauxzoon brasiliensis* in wild felids in Brazil and Argentina, and *Cytauxzoon* sp. Kozhikode in domestic cats in India (Calchi et al., 2025; Duarte et al., 2024) have been described.

The first reports of cytauxzoonosis in Europe date back to 2003, with detection of *Cytauxzoon* spp. in the Iberian lynx (*Lynx pardinus*) (Luaces et al., 2005; Meli et al., 2009; Millan et al., 2007) and in domestic cats in Spain (Criado-Fornelio et al., 2004). Subsequently, retrospective analyses revealed that *Cytauxzoon* spp. infections were already present in French wildcats as early as 1995, with a reported prevalence of 29%, and in a Swiss domestic cat in 2003 (Willi et al., 2022). Since then, *Cytauxzoon* spp. have been identified in stray and domestic cats, as well as in free-ranging and captive wild felids, across various European countries including Italy, Portugal, Spain, France, Switzerland, Germany, Hungary, and Russia (Alho et al., 2016; Antognoni et al., 2022; Carli et al., 2014; Diaz-Reganon et al., 2017; Grillini et al., 2021; Legroux et al., 2017; Naidenko et al., 2022; Nentwig et al., 2018; Panait et al., 2020; Spada et al., 2014; Veronesi et al., 2016; Willi et al., 2022). Reservoir hosts include wild felids such as the Iberian lynx, Eurasian lynx (*Lynx lynx*), and the European wildcat (*Felis silvestris*), as well as chronically infected domestic cats exhibiting prolonged parasitemia without clinical signs (Carli et al., 2012; Legroux et al., 2017).

Phylogenetic analyses based on mitochondrial gene sequences (cytochrome b and cytochrome oxidase subunit I) have revealed the presence of three distinct *Cytauxzoon* species circulating in wild felids in Europe: *Cytauxzoon europaeus* (EU1), *Cytauxzoon otrantorum* (EU2), and *Cytauxzoon banethi* (EU3) (Panait et al., 2021). Among these, *C. europaeus* seems to be most prevalent and is the only species detected so far in domestic cats in Europe. *C. europaeus* infection have been reported in wild and domestic felids in Hungary, France, Switzerland, Germany, Romania, Czeck Republic, Bosnia Herzegovina and Italy (Grillini et al., 2023; Hornok et al., 2022; Obiegala et al., 2024; Tuska-Szalay et al., 2023; Unterkofler et al., 2022; Willi et al., 2022).

European *Cytauxzoon* spp. are generally regarded as less pathogenic than *C. felis*. However, clinical disease and fatal cases have been documented in domestic cats (Alho et al., 2016; Carli et al., 2014; Carli et al., 2012; Legroux et al., 2017; Nentwig et al., 2018). Infections with *Cytauxzoon* spp. are characterized by an asexual replication in the host’s mononuclear phagocytic cells. Massive numbers of schizont-laden mononuclear cells that obstruct the vascular lumen of different organs are responsible for the severe clinical signs observed in domestic cats infected with *C. felis*. (Nietfeld and Pollock, 2002). Notably, this schizogonous stage has not been observed so far in domestic or wild felids infected with European *Cytauxzoon* spp. (Carli et al., 2014; Carli et al., 2012; Legroux et al., 2017), suggesting that the schizogonous phase in European *Cytauxzoon* spp. is probably more limited. This has also been reported for *C. felis* infections in bobcats, which often go asymptomatic (Blouin et al., 1987).

In the United States, *C. felis* is transmitted by ticks, with *Amblyomma americanum* and *Dermacentor variabilis* identified as competent vectors. Transstadial transmission has been demonstrated in these species, and ticks can transmit the parasite from both clinically ill and subclinically infected cats to susceptible hosts (Allen et al., 2019; Blouin et al., 1984; Reichard et al., 2010; Shock et al., 2011), while infected wild felids may act as reservoirs. In contrast, the tick vector responsible for the transmission of *Cytauxzoon* spp. in Europe remains unidentified. However, tick species such as *Dermacentor* spp., *Ixodes* spp., and *Rhipicephalus* spp.—all of which are present in Europe—are considered potential vectors (eCDC; Estrada Pena et al., 2017).

The aim of this study was to investigate the presence of *Cytauxzoon* spp. in ticks collected in a hotspot region of cytauxzoonosis in domestic cats in Switzerland in 2019 and in the same area during subsequent seasons. The ticks were morphologically characterized, pooled according to origin and life stage and pools investigated for the presence of *Cytauxzoon* spp. 18S rRNA by qPCR. In positive pools, 18S rRNA gene sequencing and sequencing of the mitochondrial genes (CytB and COI) was attempted.

## Materials and Methods

### Sample collection and characteristics

In February and March 2019, a total of 160 ticks were collected in a rural area close to Unterkulm, a village located in central Switzerland. A total of 62 ticks were directly collected from 7 cats from a household, in which we recently documented *C. europeaus* infections in 3 out of 10 cats (household 2 (Willi et al., 2022)). Furthermore, 98 questing ticks were collected from the vegetation at the edge of a forest located around 350 m away from this household. The questing ticks were collected by dragging a 1 m^2^ white cotton cloth over the vegetation (dragging method (Hornok et al., 2014)). An additional 505 questing ticks were collected during subsequent years from the same area (7 in September 2022, and 498 in May 2024): 423 from the edge of the forest, 33 from the meadow near the forest, and 49 from forest paths. A map from the so far detected *Cytauxzoon* spp. positive domestic and stray cats in Switzerland (adapted from (Willi et al., 2022)) and a detailed map of the region where the ticks were collected is displayed in Figure 1A and 1B respectively. Ticks were transferred to 1.5 mL Eppendorf screw tubes pre-filled with 70% ethanol and stored at room temperature until further processing. All ticks were morphologically characterized by one of the authors (R.M.E) according to published method (Eichenberger et al., 2015; Estrada-Peña, 2004; Hillyard, 1996).

**Figure 1:**
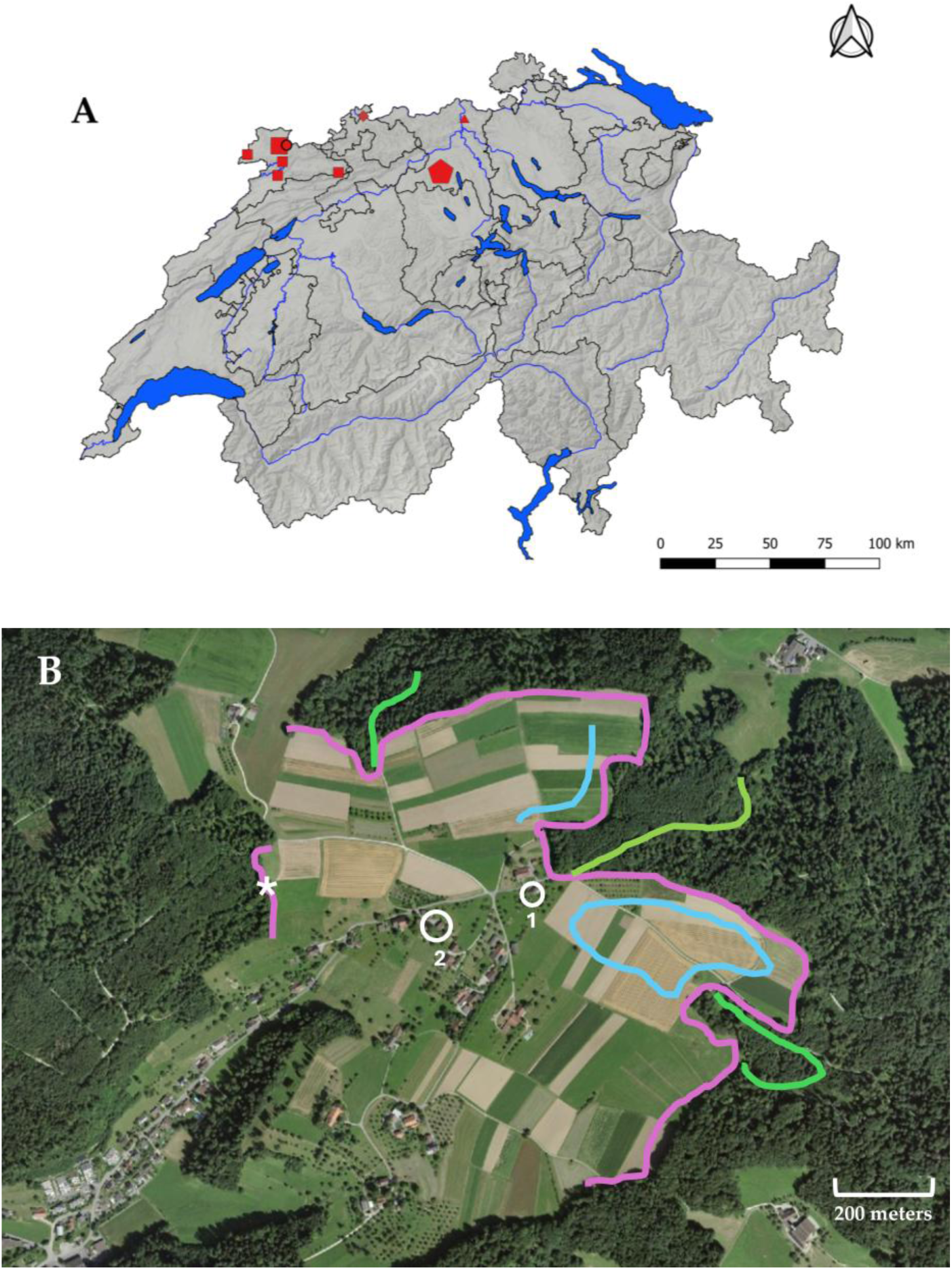
Map of Switzerland(A) showing the geographical distribution of the positive domestic and stray cat samples (Willi et al., 2022) and of the hotspot region (B) where the ticks were collected. A: The geographic origin of the *Cytauxzoon* spp positive cats from the previous study are displayed. Pentagon: region of the hotspot where the ticks were collected; circle: positive domestic cat of the Swiss-wide study 2013-2016; rhomb: positive anemic cat from the study 2019-2021; squares stray domestic cats 2014; triangle: positive domestic cat from 2003. The size of the symbols in a indicates the number of *Cytauxzoon* spp. PCR-positive or -negative samples per location. B: detailed satellite map (https://map.geo.admin.ch/) of the region of the hotspot with the two neighboring households (white circles 1 and 2 in CH 5726 Unterkulm, google plus coordinates household 2: 47.317590, 8.128228) and the regions where the ticks were collected. Pink: forest edges; green: forest paths; blue: meadow. White asterisk: region where the 3 positive questing tick pools (13, 17, 19) were collected

### Nucleic acids extraction

DNA from engorged ticks and tick pools was extracted using the QIAamp® DNA Mini Kit (Qiagen, Hombrechtikon, Switzerland) according to the manufacturer’s instructions with some modifications as previously described (Willi et al., 2007). Briefly, tick pools were pre-processed as follows: all ticks were first air-dried and then sequentially washed in 10% bleach, tap water, and distilled water. Air-dried ticks were then minced with a sterile scalpel on parafilm tape and transferred into a sterile 2 mL round-bottomed tube. A 5 mm steel bead (Retsch, Haan, Germany) and 200 mL ATL buffer were added to the minced material. The ticks were disrupted using a tissue homogenizer: either a Mixer-Mill 300 (Retsch, Haan, Germany) or a Precellys® 24 tissue homogenizer (Bertin Technologies SAS, Montigny-le-Bretonneux, France) set to 30 Hz for 1 minute. After a short centrifugation step to remove possible material from the tube lid, the steel bead was removed, and 20 mL proteinase K (Qiagen) was added, and the samples were digested overnight at 56°C. Buffer AL (400 mL) was added to the completely lysed samples and mixed thoroughly by vortexing for 15 seconds and incubated for 10 minutes at 70°C. After a short centrifugation step to remove drops from the lid, 400 μL 96% ethanol was added and mixed, and centrifuged again before loading the mixture onto the QIAamp Mini spin column (Qiagen). After the washing procedures, DNA was eluted from the column using 100 mL buffer AE and 5 minutes incubation at room temperature. DNA was stored at -80°C until further processing. At each extraction batch, a negative extraction control, consisting of 200 mL of Hank’s Balanced Salt Solution (1x HBSS, Gibco, Thermofisher Scientific, Basel, Switzerland) was run in parallel.

### Diagnostic assays, amplification of the 18S rRNA, CytB and COI genes and sequencing

Quality and quantity of the DNA samples extracted from ticks were tested with a commercial 18S rRNA real-time TaqMan qPCR assay (Thermofisher Scientific) as described previously (Boretti et al., 2009). Only samples with cycle threshold (Ct) values < 30 were further analyzed and screened for *Cytauxzoon spp.* using a real-time TaqMan qPCR assay that amplifies 69 bp of the 18S rRNA gene (Meli et al., 2009; Nentwig et al., 2018; Willi et al., 2022). Assays were run on an ABI 7500 Fast Real-Time PCR system (Applied Biosystems, Rotkreuz, Switzerland). Positive and negative PCR controls were run with each PCR assay and consisted of DNA from a *Cytauxzoon spp.* PCR-positive Iberian lynx (confirmed by sequencing) and nuclease-free water, respectively. All samples with threshold cycle (Ct) values < 35 were subjected to confirmation by a conventional PCR that amplifies 221 bp of the 18S rRNA gene of *Cytauxzoon spp*. (Meli et al., 2009; Nentwig et al., 2018).

Samples that were PCR positive in the 18S rRNA conventional PCR underwent amplification and sequencing of cytochrome b (CytB) and cytochrome oxidase subunit I (COI) mitochondrial genes of *Cytauxzoon spp.*, either directly or after a pre-amplification step using the Prelude™ PreAmp Master Mix (TaKaRa Bio, Kusatsu, Japan), as previously described (Willi et al., 2022). The PCR products were separated on a 2% agarose gel, and bands of appropriate size were sequenced at a commercial laboratory (Microsynth AG, Balgach, Switzerland) using the amplification primers. Sequences were edited and assembled using Geneious Prime® 2020.2.5 software (https://www.geneious.com; Biomatters Limited, Auckland, New Zealand) (Kearse et al., 2012). Sequence identification was conducted by comparing the obtained sequences to existing sequences through the BLASTn search program (http://www.ncbi.nlm.nih.gov/blast/Blast.cgi).

Nucleotide sequences obtained in this study, with exception of the sequence from pool 19 (too short for the submission, < 200 bp) have been submitted to GenBank under accession numbers PV944017 (tick pool 7), PV944018 (tick pool 13), PV944019 (tick pool 17).

## Results

### Ticks collected in the hotspot region in 2019

A total of 160 ticks collected in 2019 were analyzed. All ticks belonged to the species *Ixodes ricinus* and comprised 14 males, 49 females, and 97 nymphs. They were allocated to 42 pools according to origin, sex, and developmental stage (Table 1). Sixty-two ticks were collected from domestic cats residing in one of the households in which *C. europaeus* infection was documented in several cats (Willi et al., 2022). An additional 98 ticks (2 females and 96 nymphs) were collected from vegetation at the forest edge, approximately 350 meters northwest of the household.

**Table 1.**
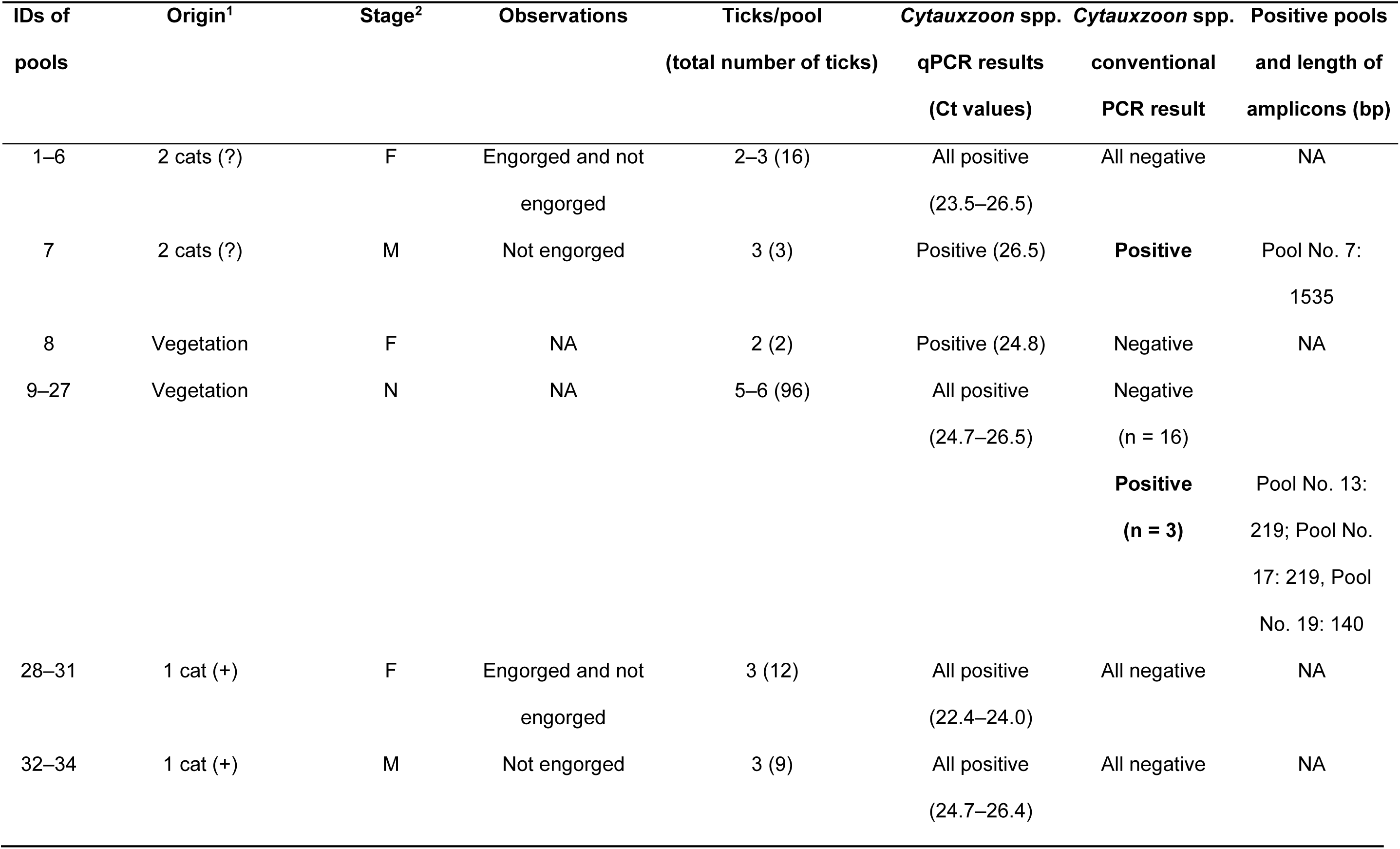
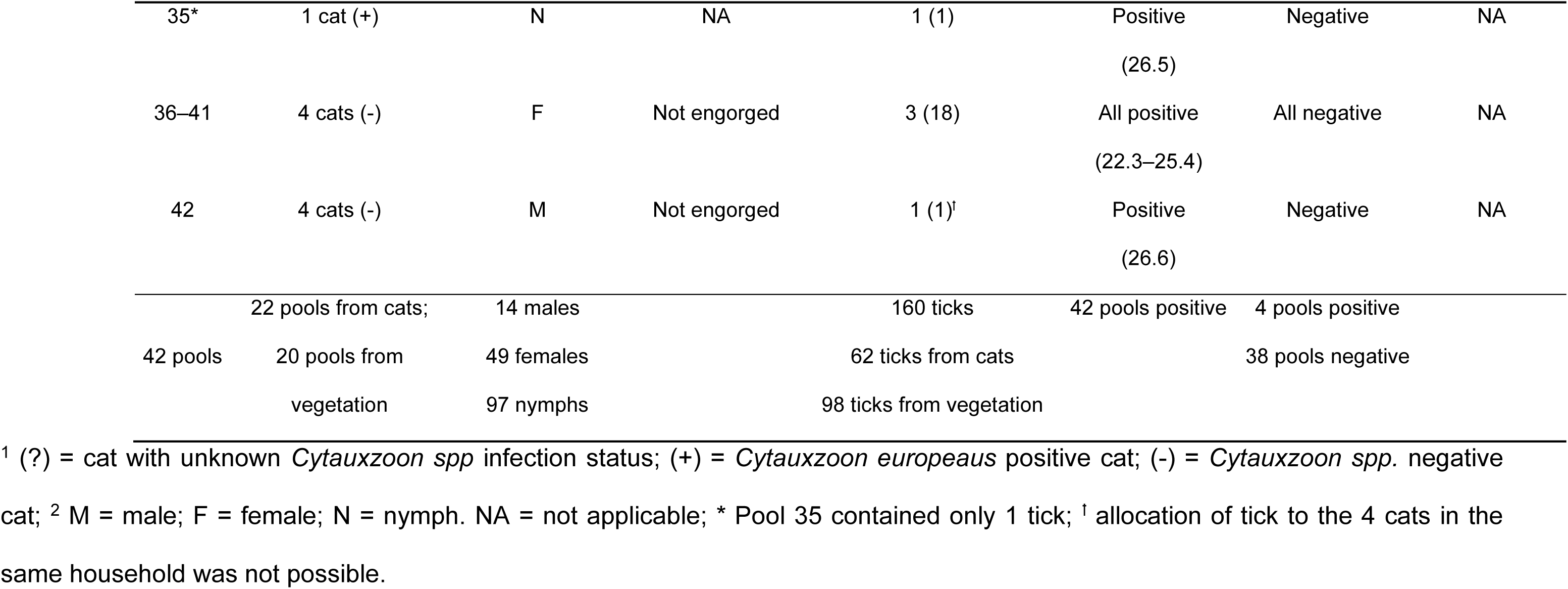
Characteristics of the investigated *I. ricinus* tick pools collected in 2019 and *Cytauxzoon spp*. PCR results.

All 42 tick pools tested positive for *Cytauxzoon* spp. using qPCR screening and were subsequently analyzed by conventional PCR targeting a 221 bp fragment of the 18S rRNA gene. Of these, four pools yielded weakly positive amplicons. Sequencing confirmed the presence of *Cytauxzoon* spp. in these four pools, with three yielding partial sequences of 140–219 bp and one yielding an almost full-length sequence of 1535 bp of the 18S rRNA gene (Table 1).

The *Cytauxzoon*-positive pools included one pool (No. 7) containing four non-engorged male *Ixodes ricinus* ticks collected from two domestic cats of unknown *Cytauxzoon* spp. infection status, and three pools (Nos. 13, 17, and 19) containing 5-6 *I. ricinus nymphs* each collected from vegetation.

BLAST analysis of the 1535 bp sequence of pool No. 7 revealed 100% identity with *Cytauxzoon* spp. sequences previously obtained from two domestic cats in the neighboring household, as well as from two French wildcats sampled in 1995 (Willi et al., 2022). Additionally, this sequence was identical to those of *Cytauxzoon* spp. isolated from two Italian domestic cats collected in 2016/2017 (unpublished dataset, GenBank accession numbers OM004051 and OM004053).

The partial sequences from tick pools No. 13, 17, and 19 showed 98.6%, 99.5%, and 99.3% identity, respectively, to *Cytauxzoon* spp. sequences—including *Cytauxzoon europaeus*—previously detected in European wild felids (Obiegala et al., 2024; Panait et al., 2021; Unterkofler et al., 2022; Willi et al., 2022) and Swiss domestic cats (Nentwig et al., 2018; Willi et al., 2022).

All attempts to amplify and sequence the mitochondrial genes (CytB and COI) from the positive pools were unsuccessful.

### Ticks collected in 2022 and 2024

In 2022 and 2024–approximately three and a half and five years after the first collection in the endemic region (Willi et al., 2022)–an additional 505 ticks were collected from the same area and the ticks analyzed for the presence of *Cytauxzoon* spp.

All 505 ticks belonged to the species *I. ricinus* and comprised 36 males, 34 females, and 428 nymphs. They were allocated to 143 pools according to origin, sex, and developmental stage (Table 2).

**Table 2.**
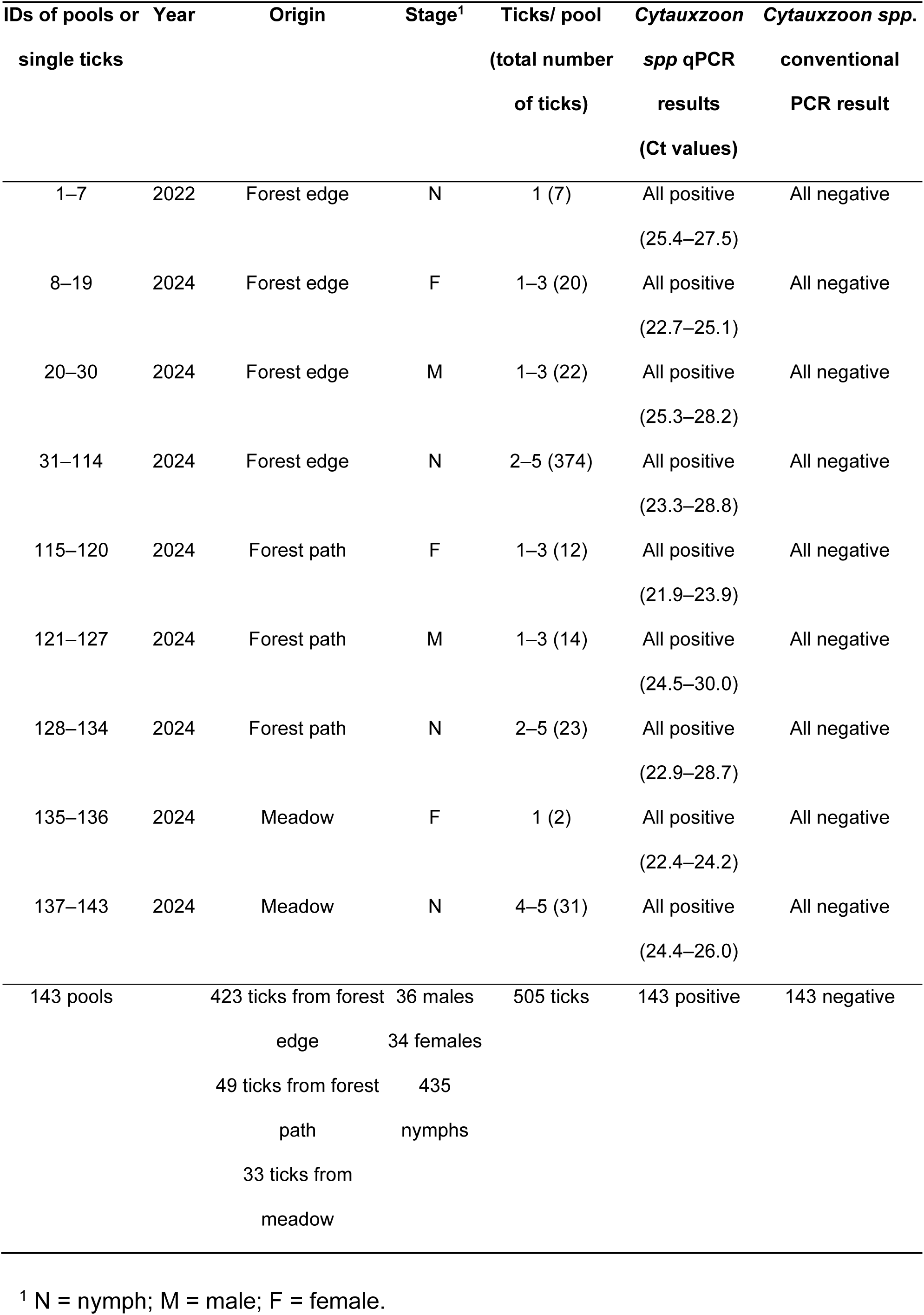
Characteristics of the investigated questing *I. ricinus* tick pools collected in 2022 and 2024 and *Cytauxzoon spp*. PCR results.

| IDs of pools or<br>single ticks | Year | Origin | Stage<sup>1</sup> | Ticks/ pool<br>(total number<br>of ticks) | *Cytauxzoon*<br>*spp* qPCR<br>results<br>(Ct values) | *Cytauxzoon spp.*<br>conventional<br>PCR result |
| --- | --- | --- | --- | --- | --- | --- |
| 1–7 | 2022 | Forest edge | N | 1 (7) | All positive<br>(25.4–27.5) | All negative |
| 8–19 | 2024 | Forest edge | F | 1–3 (20) | All positive<br>(22.7–25.1) | All negative |
| 20–30 | 2024 | Forest edge | M | 1–3 (22) | All positive<br>(25.3–28.2) | All negative |
| 31–114 | 2024 | Forest edge | N | 2–5 (374) | All positive<br>(23.3–28.8) | All negative |
| 115–120 | 2024 | Forest path | F | 1–3 (12) | All positive<br>(21.9–23.9) | All negative |
| 121–127 | 2024 | Forest path | M | 1–3 (14) | All positive<br>(24.5–30.0) | All negative |
| 128–134 | 2024 | Forest path | N | 2–5 (23) | All positive<br>(22.9–28.7) | All negative |
| 135–136 | 2024 | Meadow | F | 1 (2) | All positive<br>(22.4–24.2) | All negative |
| 137–143 | 2024 | Meadow | N | 4–5 (31) | All positive<br>(24.4–26.0) | All negative |
| 143 pools |  | 423 ticks from forest<br>edge<br>49 ticks from forest<br>path<br>33 ticks from<br>meadow | 36 males<br>34 females<br>435<br>nymphs | 505 ticks | 143 positive | 143 negative |

<sup>1</sup> N = nymph; M = male; F = female.

All 143 tick pools tested positive for *Cytauxzoon spp.* using qPCR screening and were subsequently analyzed by conventional PCR targeting a 221 bp fragment of the 18S rRNA gene. However, none of the individual ticks or pooled samples yielded a positive result in the conventional PCR assay (Table 2).

## Discussion

This study reports, for the first time, the detection of *Cytauxzoon* spp. in questing *I. ricinus* ticks. To date, the tick vector for European *Cytauxzoon* spp. is unknown. Our data suggests that *Ixodes ricinus* might be involved in the transmission cycle of this pathogen in Central Europe. *Cytauxzoon* spp. was detected in ticks collected from cats living in households with *Cytauxzoon* spp. infected cats, but also in questing ticks collected in an area located around 350 meters (∼380 yards) away from this household. A pool of two unengorged male *I. ricinus* ticks obtained from two cats with unknown *Cytauxzoon* infection status, as well as three pools of *I. ricinus* nymphs collected directly from vegetation in the same area, tested positive by conventional PCR. Subsequent sequencing of a partial fragment of the 18S rRNA gene confirmed the taxonomic assignment to the *Cytauxzoon* genus. Based on the 18S rRNA gene sequence, the presence of *Cytauxzoon* spp. could be confirmed in the questing ticks; however, species-level identification was not possible, likely due to the low parasitic load in the tick samples, despite DNA preamplification. Given that only *C. europaeus* has been reported in domestic and wild felids in Switzerland (Willi et al., 2022), and that the positive ticks were collected concurrently with the occurrence of *C. europaeus* infections in several cats of the household, the detected organisms in the ticks likely represent *C. europaeus*. However, we were not able to amplify and sequence the mitochondrial genes COI and CytB of *Cytauxzoon* spp. from the positive tick pools, which would be a prerequisite to definitely assign the amplified species to *C. europeaus*. Therefore, a second attempt was undertaken in autumn 2022 and spring 2024 to collect more questing ticks from the same area, where cytauxzoonosis was considered to be endemic (Willi et al., 2022). However, no further *Cytauxzoon*-positive ticks were identified at that time. Given that infected cats typically remain PCR-positive for *Cytauxzoon* spp. for years, even after antibiotic treatment (Nentwig et al., 2018), it was expected that these chronically infected animals with prolonged parasitemia would contribute to a sustained parasitic burden within the local tick population. However, possibly increased implementation of tick prophylaxis measures—potentially prompted by the earlier *Cytauxzoonosis* infection—may have significantly reduced tick exposure in domestic animals, leading to a subsequent decrease in tick infestation rates and, consequently, a lower prevalence of *Cytauxzoon* spp. in the tick population.

For *C. felis*, *D. variabilis* and *A. americanum* ticks have been shown to be competent tick vectors (Blouin et al., 1984; Reichard et al., 2010). Neither tick species is present in Central Europe (Estrada Pena et al., 2017). *Ixodes ricinus* is by far the most common tick species in Switzerland and is present in most parts of the country (Aeschlimann, 1972), but also highly prevalent in other central European countries (Estrada Pena et al., 2017). *Ixodes ricinus* ticks are known vectors of many pathogens of veterinary relevance, among them other piroplasms like *Babesia* (i.e., *Babesia divergens*, *Babesia microti,* and *Babesia venatorum* (Bonnet et al., 2009; Bonnet et al., 2007; Gray et al., 2002)) as well as *Theileria* spp. (Fernandez et al., 2022). *Babesia* spp. are phylogenetically related to *Cytauxzoon* spp., but they differ in their life cycle compared to *C. felis*, since *Babesia* spp. do not undergo a schizogenous stage. However, schizogonoy has not yet been documented in domestic and wild felids infected with European *Cytauxzoon* spp. (Willi et al., 2022). Besides *I. ricinus*, *Dermacentor marginatus*, *Dermacentor reticulatus*, *Haemaphysalis punctata*, *Rhipicephalus sanguineus,* and other species of the genus *Ixodes* spp. (*Ixodes hexagonus*) have also been described in Switzerland (Aeschlimann and Papadopoulos, 1998; Bernasconi et al., 2002; Bernasconi et al., 1997), and *H. punctata* has been shown to transmit *Babesia major* (Phipps et al., 2022).

In this study, a total of 603 questing *I. ricinus* ticks were collected, with the majority being nymphs (88%), and only a smaller proportion consisting of females and males. We used dragging, which is, along with flagging, a commonly used method for collecting questing ticks, including *I. ricinus*. A study comparing both methods found that flagging was generally more efficient in collecting adult ticks, especially in spring and winter (Dantas-Torres et al., 2013). This may explain the large predominance of nymphs found in our study. Both methods have also been successfully used to collect other tick species, such as *D. marginatus*, *Hyalomma marginatum,* and *Haemaphysalis inermis.* However, these species are not present in central Switzerland, explaining why we exclusively found *I. ricinus*.

The majority of tick samples were collected in the vegetation at the border between the forest and meadow. Fewer ticks were collected along forest paths and in the meadow. *Ixodes ricinus* ticks thrive in habitats that provide high humidity, mild temperatures, dense vegetation, and suitable hosts. Areas with dense shrub layers and transitional zones between forests and open lands (ecotones), such as the forest edges and paths, provide ideal conditions. These habitats not only retain moisture but also attract a variety of wildlife, increasing tick-host interactions (Estrada-Pena, 2001). The tick abundance observed in our study appears to reflect these habitat characteristics, with meadows and forest path edges likely supporting lower tick densities due to reduced humidity retention, as these areas typically lack the leaf litter necessary to maintain stable microclimatic moisture conditions.

The detection of *Cytauxzoon spp.* in questing ticks collected from the vegetation speaks against the possibility that the uptake of blood from an infected cat caused the PCR-positive signal, but points towards a life cycle for *Cytauxzoon* spp. in this tick species. Particularly, the detection in nymphs is noteworthy as it addresses the possibility of transovarial transmission from an infected female tick to her offspring without prior host feeding. This transmission mode would have important implications for the persistence and spread of the pathogen in the *I. ricinus* population (Hauck et al., 2020). Transovarial transmission has been confirmed for *Babesia canis* in *D. reticulatus* ticks in Europe (Mierzejewska et al., 2018), but it has not yet been investigated in *Cytauxzoon* spp., including *C. felis* (Wikander and Reif, 2023). Although *Cytauxzoon* spp. PCR-positive nymphs were detected in 2019; the absence of positive ticks in follow-up surveys in the same area several years later raises doubts about the occurrence of transovarial transmission. However, it should be noted that the ticks in later years were collected near, but not at, the exact location where the PCR-positive nymphs were originally detected. Moreover, the extent of general tick infestation with *Cytauxzoon* spp. in 2019 in the area remains unknown, and a depletion of the transmission cycle in case of strict tick prevention in cats might have happened. Thus, further investigations are required to clarify whether and to what extent *Cytauxzoon* spp. can be vertically transmitted in *I. ricinus*.

The *Cytauxzoon* spp. qPCR assay used to screen tick DNA for the presence of the hemoparasite demonstrated high sensitivity when applied to blood samples from felids (Nentwig et al., 2018; Willi et al., 2022). Moreover, *in silico,* the assay showed high specificity for detecting *Cytauxzoon* spp., which was confirmed using blood samples from infected cats (Willi et al., 2022). However, when applied to investigating ticks, this same assay yielded positive results for all tested samples. Each sample pool underwent additional verification using conventional PCR targeting a segment of the 18S rRNA gene of *Cytauxzoon* spp. to confirm the presence of parasite-specific sequences in the tick pools, and many of them tested negative. Moreover, there was no correlation between the Ct value of the qPCR and the results in the conventional PCR. The qPCR Ct values from ticks collected in 2022 and 2024 were comparable to those from the samples collected in 2019. At the same time, some of the 2019 samples tested positive, but all of the 2022/2024 samples tested negative by conventional PCR. We therefore hypothesize that the qPCR lacks specificity when applied to tick DNA, as it co-amplifies DNA from tick endosymbionts. Ticks host a diverse array of endosymbiotic bacteria, which are crucial for their survival and reproduction and may play a possible role in the transmission dynamics of tick-borne diseases (Greay et al., 2018; Wiesinger et al., 2023). Therefore, the potential for cross-amplification of endosymbiont DNA in qPCR assays targeting the *Cytauxzoon* spp. 18S rDNA gene necessitates careful interpretation and confirmation of results, particularly when analyzing tick-derived DNA.

In conclusion, cytauxzoonosis is a significant vector-borne disease of felids, with potentially serious health consequences for infected animals. While substantial progress has been made in recent years regarding the presence and geographic distribution of various *Cytauxzoon* spp. in Europe as well as worldwide, information on the associated tick vectors remains limited. This study provides the first evidence that *I. ricinus* may be a potential vector for European *Cytauxzoon* spp. and suggests the possibility of transovarial transmission of these pathogens in *I. ricinus*. These findings provide a foundation for future epidemiological studies aimed at confirming vector competence, clarifying transmission dynamics, and determining whether *I. ricinus* or other tick species in Europe play a role in the life cycle of *Cytauxzoon* spp.

## Acknowledgements

The authors thank L. Janowitz, and E. Pestana Mesquita for their excellent assistance in sample collection. The laboratory work was performed using the logistics of the Center for Clinical Studies, Vetsuisse Faculty, University of Zurich.

## Notes

### Competing Interest Statement

The authors have declared no competing interest.

